# AdmixPipe: Population analyses in Admixture for non-model organisms

**DOI:** 10.1101/2020.07.06.190389

**Authors:** Steven M. Mussmann, Marlis R. Douglas, Tyler K. Chafin, Michael E. Douglas

**Author notes:** Corresponding author and person to whom reprint requests should be addressed: Steven M. Mussmann, Department of Biological Sciences, University of Arkansas, Fayetteville, AR 72701, Voice.

## Abstract

**Background:** Research on the molecular ecology of non-model organisms, while previously constrained, has now been greatly facilitated by the advent of reduced-representation sequencing protocols. However, tools that allow these large datasets to be efficiently parsed are often lacking, or if indeed available, then limited by the necessity of a comparable reference genome as an adjunct. This, of course, can be difficult when working with non-model organisms. Fortunately, pipelines are currently available that avoid this prerequisite, thus allowing data to be *a priori* parsed. An oft-used molecular ecology program (i.e., Structure), for example, is facilitated by such pipelines, yet they are surprisingly absent for a second program that is similarly popular and computationally more efficient (i.e., Admixture). The two programs differ in that Admixture employs a maximum-likelihood framework whereas Structure uses a Bayesian approach, yet both produce similar results. Given these issues, there is an overriding (and recognized) need among researchers in molecular ecology for bioinformatic software that will not only condense output from replicated Admixture runs, but also infer from these data the optimal number of population clusters (K).

**Results:** Here we provide such a program (i.e., AdmixPipe) that (a) filters SNPs to allow the delineation of population structure in Admixture, then (b) parses the output for summarization and graphical representation via Clumpak. Our benchmarks effectively demonstrate how efficient the pipeline is for processing large, non-model datasets generated via double digest restriction-site associated DNA sequencing (ddRAD). Outputs not only parallel those from Structure, but also visualize the variation among individual Admixture runs, so as to facilitate selection of the most appropriate *K*-value.

**Conclusions:** AdmixPipe successfully integrates Admixture analysis with popular variant call format (VCF) filtering software to yield file types readily analyzed by Clumpak. Large population genomic datasets derived from non-model organisms are efficiently analyzed via the parallel-processing capabilities of Admixture. AdmixPipe is distributed under the GNU Public License and freely available for Mac OSX and Linux platforms at: https://github.com/stevemussmann/admixturePipeline.

## Background

Advances in genomics during the past decade have accelerated research in molecular ecology by significantly increasing the capacity of researchers to generate vast quantities of data at relatively low cost. These advances largely represent the development of reduced representation genomic libraries [1–3] that identify tens of thousands of SNPs for non-model organisms, coupled with high-throughput sequencing methods that efficiently genotype fewer SNPs for thousands of individuals [4]. However, data generation, particularly through these novel and affordable marker-discovery methods [5], has greatly outpaced analytical capabilities, and especially so with regard to evolutionary and conservation genomics.

Here, technological advances have also precipitated a suite of new analytical issues. The thousands of SNPs generated in a typical RADseq project may exhibit biases that impact the inferences that can be drawn from these data [6], and which necessitate careful data filtration to avoid [7]. Yet, the manner by which data are filtered represents a double-edged sword. While it is certainly mandated (as above), the procedures involved must be carefully evaluated in the context of each study, in that downstream analyses can be seriously impacted [8, 9], to include the derivation of population structure [10].

For example, the analysis of multilocus codominant markers in evaluation of population structure is frequently accomplished using methods that make no *a priori* assumptions about underlying structure. One of the most popular options to accomplish this is the program Structure [11–13]. However, it necessitates that users test specific clustering values (K), and conduct *post hoc* evaluation of these results so as to determine an optimal K [14]. This typically involves searching a complicated parameter space using heuristic algorithms for Maximum Likelihood (ML) and Bayesian (BA) methods that, in turn, provide additional complications such as a tendency to sample local optima [15].

A common strategy to mitigate this is to sample multiple independent replicates at each K, using different random number seeds for initialization. These results are subsequently collated and evaluated to assess confidence that global rather than local optima have indeed been sampled. Clearly, this procedure must be automated so as to alleviate the onerous task of testing multiple replicates across a range of K-values. Pipelines to do so are available for Structure, and have been deployed on high-performance computing systems via integrated parallelization (StrAuto, ParallelStructure) [16, 17]. Multiple programs have likewise been developed for handling Structure output (i.e., Clumpp, Distruct) [18, 19]; and pipelines constructed to assess the most appropriate K-values (i.e., StructureHarvester, Clumpak) [20, 21].

Despite the considerable focus on Structure, few such resources have been developed for a popular alternative program (i.e., Admixture [22]). The Web of Science indexing service indicates that (as of January, 2020) it has been cited 1,812 times since initial publication (September, 2009). This includes 479 (26.4%) in 2019 alone. Despite its popularity, it has just a single option that promotes the program as part of a pipeline (i.e., SNiPlay3 [23]), and unfortunately it requires a reference genome as an adjunct for its application. Needless to say, its applicability is thus limited for those laboratories that employ non-model organisms as study species.

Options for post-processing of Admixture results are similarly deficit. However, one positive is that Clumpak is flexible enough in its implementation to allow for the incorporation of Admixture output, as well as that of Structure. Furthermore, no available software currently exists that can summarize the variation in cross-validation (CV) values, the preferred method for selecting an optimal K-value in Admixture [24].

Here we describe a novel software package that integrates Admixture as the primary component of an analytical pipeline that also incorporates the filtering of data as part of its procedure. This, in turn, provides a high-throughput capability that not only generates input for Admixture but also evaluates the impact of filtering on population structure. AdmixPipe also automates the process of testing multiple K-values, conducts replicates at each K, and automatically formats these results as input for the Clumpak pipeline. Optional post-processing scripts are also provided as a part of the toolkit to process Clumpak output, and to visualize the variability among CV values for independent Admixture runs. Sections of the pipeline are specifically designed for use with non-model organisms, as these are increasingly common study species in evolutionary and conservation genomic investigations.

## Implementation

AdmixPipe requires two input files: a population map and a standard VCF file. The population map is a tab-delimited text file with each row representing a sample name/ population pair. The VCF file is filtered according to user-specified command line options that include the following: minor allele frequency (MAF) filter, biallelic filtering, data thinning measured in basepairs (bp), and missing data filtering (for both individuals and loci). Users may also remove specific samples from their analysis by specifying a file of sample names to be ignored. All filtering and the initial conversion to Plink (PED/MAP) format [25] is handled by VCFTOOLS [26].

AdmixPipe is intended for use with non-model organisms that lack genomic reference data, and given this, additional conversions are required before the Plink-formatted files will be accepted by Admixture. Popular software packages for *de novo* assembly of RADseq data, such as pyRAD [27, 28] produce VCF files with each locus as an individual “chromosome.” This, in turn, yields output that exceeds the number of chromosomes in those model organisms for which Plink was originally designed. The initial MAP file is therefore modified to append a letter at the start of each “chromosome” number. Plink is then executed using the “–allow-extra-chr 0” option that treats loci as unplaced contigs in the final PED/MAP files submitted to Admixture.

The main element of the pipeline executes Admixture on the filtered data. The assessment of multiple K values and multiple replicates is automated based upon user-specified command line input. The user defines minimum and maximum K values to be tested, in addition to the number of replicates for each K. Users may also specify the number of processor cores to be utilized by Admixture, and the cross-validation number which is utilized in determining optimal K. The final outputs of the pipeline include a compressed results file and a population file that are submitted as-is to Clumpak for processing and visualization.

The pipeline also offers two accessory scripts for processing of Clumpak output. The first (i.e., distructRerun.py) compiles the major clusters identified by Clumpak, generates Distruct input files, executes Distruct, and extracts CV-values for all major cluster runs. The second script (i.e., cvSum.py) plots the boxplots of CV-values against each K so as to summarize the distribution of CV-values for multiple Admixture runs. This permits the user to make an informed decision on the optimal K by graphing how these values vary according to independent Admixture runs.

Admixture is the only component of the pipeline that is natively parallelized. Therefore, we performed benchmarking to confirm that processing steps did not significantly increase runtime relative to that expected for Admixture. Data for benchmarking were selected from a recently published paper that utilized AdmixPipe for data processing [29]. The test data contained 343 individuals and 61,910 SNPs. Four data thinning intervals (i.e.,1, 25, 50, and 100) yielded SNP datasets of variable size for performance testing. All filtering intervals were repeated with variable numbers of processor cores (i.e.,1, 2, 4, 8, and 16). Sixteen replicates of Admixture were first conducted for each K=1-8 at each combination of thinning interval and number of processor cores, for a total of 20 executions of the pipeline. The process was then repeated for each K=9-16, for an additional 20 runs of the pipeline. Memory profiling was conducted through the python3 ‘mprof’ package at K=16, with a thinning interval of 1 as a final test of performance. All tests were completed on a computer equipped with dual Intel Xeon E5-4627 3.30GHz processors, 256GB RAM, and with a 64-bit Linux environment.

## Results

The filtering intervals resulted in datasets containing 61,910 (interval = 1bp), 25,851 (interval = 25bp), 19,140 (interval = 50bp), and 12,527 SNPs (interval = 100bp). Runtime increased linearly with the number of SNPs analyzed, regardless of the number of processors utilized (Figure 1: R^2^ = 0.975, df = 58). For example, increasing the number of SNPs from 12,527 to 61,910 (494% increase) produced an average increase of 519% in AdmixPipe runtime (SD = 41.6%).

**Figure 1.**
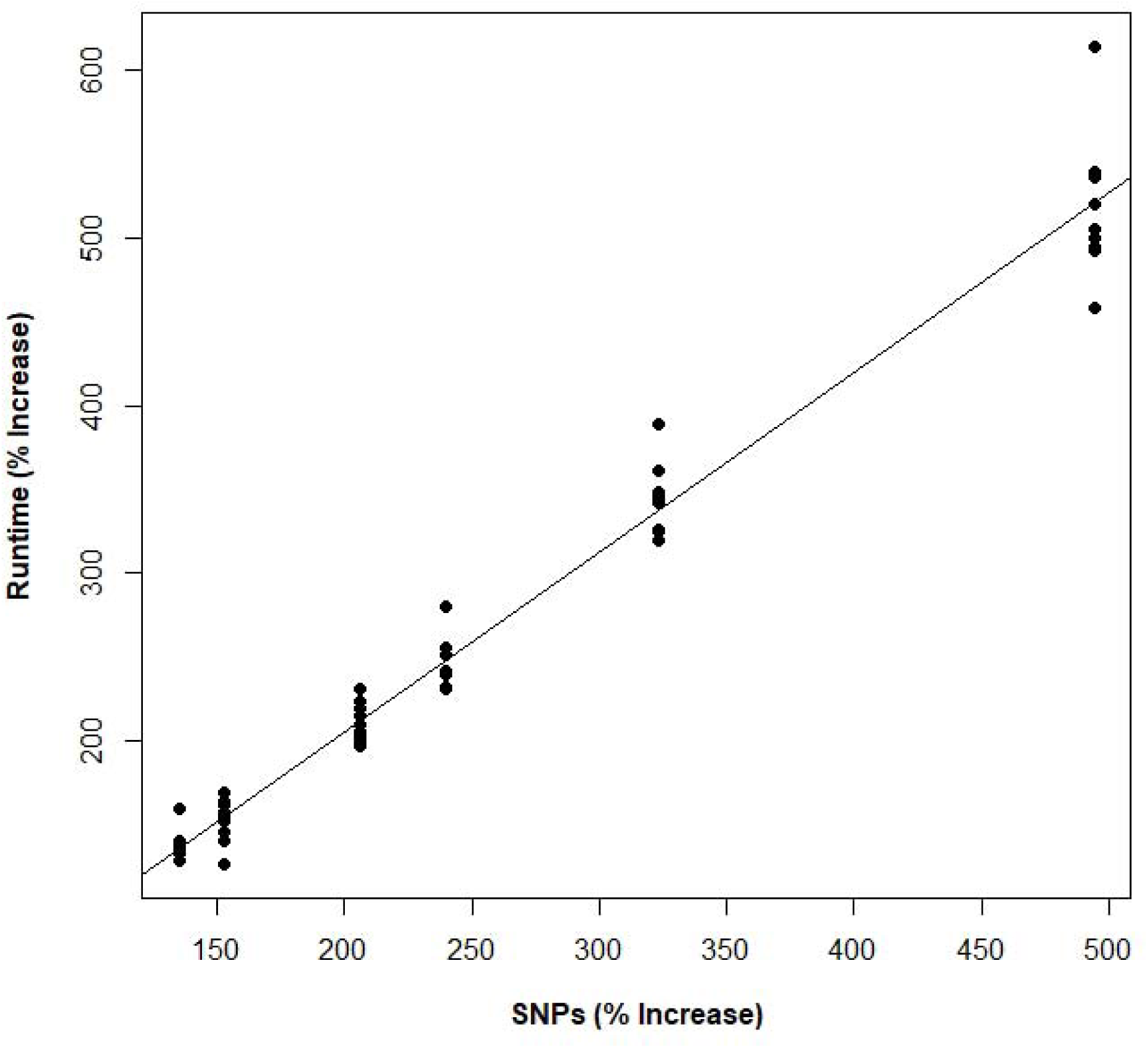
The percent increase in runtime for AdmixPipe exhibits a nearly 1:1 ratio with respect to percent increase in the number of SNPs. Data is based upon pairwise comparisons in runtime and input size increases for four datasets of varying size (61,910 SNPs, 25,851 SNPs,, 19,140 SNPs, and 12,527 SNPs). R^2^ = 0.975, degrees of freedom=58.

Little change was observed in response to increasing the numbers of processor cores from K=1-8 (Figure 2A). A slight decrease in performance was observed in some cases, particularly for the largest dataset. This trend changed at higher K-values, as substantial gains were observed at K=9-16 when processors were increased from 1 to 4. The most dramatic performance increase was observed for the 61,910 SNP dataset, where a 24.3-hour (34.5%) reduction in computation time occurred when processors increased from 1 to 4. However, only marginal improvements occurred when processors were increased from 1 to 8 (24.5 hours; 34.7%) or 16 (26.2 hours; 37.7%).

**Figure 2.**
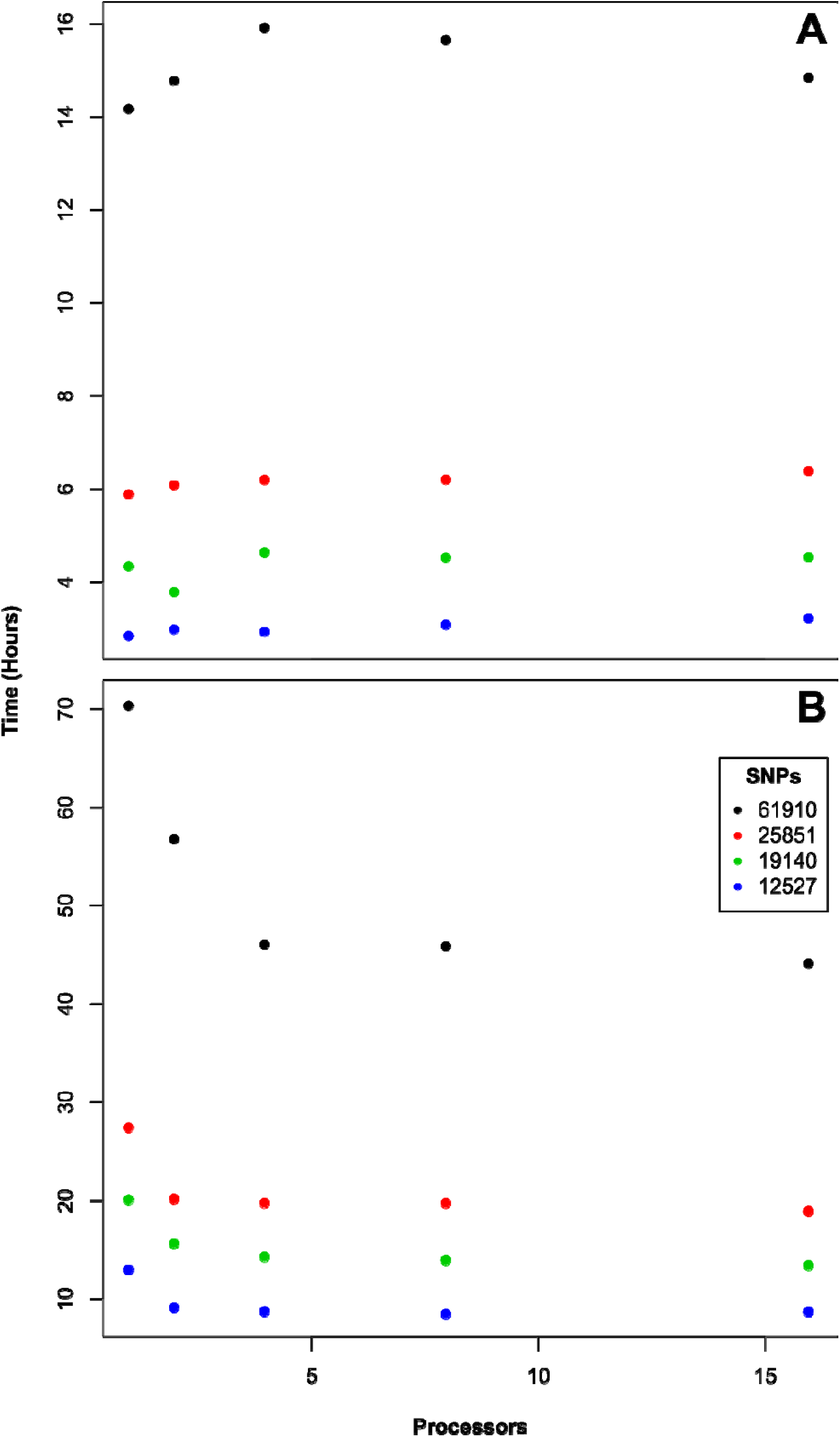
Results of AdmixPipe for two ranges of population clustering (K) values. Time is presented in hours on the Y-axis. Plot A shows total runtime for 20 replicates each of K=1-8. Plot B shows total runtime for 16 replicates each of K=9-16. The number of processor cores (CPU=1, 2, 4, 8, and 16) was varied across runs. Four data thinning intervals (1, 25, 50, and 100) produced variable numbers of SNPs (61,910, 25,851, 19,140, and 12,527 respectively).

Profiling also revealed efficient and consistent memory usage. The greatest memory spike occurred during the initial filtering steps, when peak memory usage reached approximately 120 MB. All subsequent usage held constant at ∼60 MB as Admixture runs progressed.

## Discussion

The performance of AdmixPipe improved with the number of processor cores utilized at higher K-values. However, it did not scale at the rate suggested in the original Admixture publication. We have been unable to attribute the difference in performance to any inherent property of our pipeline. Filtering and file conversion steps at the initiation of AdmixPipe are non-parallel sections. Reported times for completion of these steps were approximately constant across runs, with the maximum reported time being eight seconds. This indicates that Admixture itself is the main driver of performance, as it comprises the vast majority of system calls made by AdmixPipe.

The original performance increase documented for Admixture was 392% at K=3, utilizing four processor cores [24]. Unfortunately, we could not replicate this result with our benchmarking data [29], or the original test data (i.e.,324 samples; 13,928 SNPs) [24] which parallels our own. When we attempted to replicate the original benchmark scores, we found that it also failed to scale as the number of processor cores increased (1-core 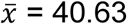 seconds, *σ* = 0.90; 4-core 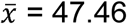 seconds, *σ* = 4.71). Furthermore, we verified that performance did increase with up to four processor cores at higher K values (K≥9). We therefore view this as ‘expected behavior’ for Admixture, and find no reason to believe that AdmixPipe has negatively impacted the performance of any individual program.

Results of AdmixPipe were similar to those found by Structure for the test dataset, as evaluated in an earlier publication [29], and gauged for the optimum K=8. This is not surprising, given that Admixture implements the same likelihood model as does Structure [22]. However, minor differences have previously been noted for both programs in the assignment probabilities [29, 30].

Memory usage was efficient and constant, with the greatest increase occurring when Plink was executed. Thus, users will be able to execute AdmixPipe on their desktop machines for datasets sized similarly to that evaluated herein. Performance gains were minimal with >4 processors, and this (again) reduces the necessity for supercomputer access, since desktop computers with ≥4 processor cores are now commonplace. However, given the built-in parallelization capabilities of AdmixPipe, its application on dedicated high-performance computing clusters will be beneficial when runtime considerations are necessary, such as when evaluating K>8, or SNPs≥20,000.

Finally, our integration of common SNP filtering options provides the flexibility to quickly filter data and assess the manner by which various filtering decisions impact results. A byproduct of the filtering process is the production of a Structure-formatted file that will facilitate comparisons with other popular algorithms that assess population structure. These options are important tools, particularly given recent documentation regarding of the impacts of filtering on downstream analyses. We thus suggest that users implement existing recommendations on filtering RAD data, and use these to investigate subsequent impacts on their own data [7–10].

## Conclusions

Benchmarking has demonstrated that the benefits of AdmixPipe (e.g., low memory usage and performance scaling with low numbers of processor cores at high K-values) will prove useful for researchers with limited access to advanced computing resources. AdmixPipe also allows the effects of common filtering options to be assessed on population structure of study species by coupling this process with the determination of population structure. Integration with Clumpak, and our custom options that allow plotting of data, to include variability in CV-values and customization of population-assignment plots, will facilitate the selection of appropriate K-values and allow variability to be assessed across runs. These benefits thus allow researchers to implement recommendations regarding assignment of population structure in their studies, and to accurately report the variability found in their results [31]. In conclusion, AdmixPipe is a new tool that successfully fills a contemporary gap found in pipelines that assess population structure. It is our hope that AdmixPipe, and its subsequent improvements will greatly facilitate the analysis of SNP data in non-model organisms.

## Acknowledgements

Computational resources were provided by the Arkansas High Performance Computing Center (AHPCC) and the NSF Jetstream XSEDE Resource (XSEDE Allocation: TG-BIO160065). This research represents partial fulfillment of the Ph.D. degree (SMM) in Biological Sciences at University of Arkansas.

## Availability of data and materials

Data utilized for benchmarking was part of an earlier publication, and is available on Data Dryad (https://datadryad.org/stash/dataset/doi:10.5061/dryad.d3q3220). Source code for AdmixPipe is released under the GNU General Public License v3.0 at https://github.com/stevemussmann/admixturePipeline. The pipeline will run on Unix-based operating systems such as Mac OSX and Linux. It is compatible with Python 2.7+ and Python 3.5+. Dependencies include other freely available software packages (Admixture, Distruct, Plink, and VCFtools).

## Authors’ Contributions

SMM, MRD, and MED designed the study; SMM and TKC authored the Python code for AdmixPipe; TKC and SMM completed data analyses and program testing; all authors contributed in drafting the manuscript, and all approved the final version.

## Consent for publication

Not applicable.

## Ethics approval and consent to participate

Not applicable.

